# Verifying the concordance between motion corrected and conventional MPRAGE for pediatric morphometric analysis

**DOI:** 10.1101/2024.10.01.616016

**Authors:** Barat Gal-Er, Yannick Brackenier, Alexandra F. Bonthrone, Chiara Casella, Anthony Price, Sophie Arulkumaran, Andrew T.M. Chew, Chiara Nosarti, Michela Cleri, Pierluigi Di Cio, Alexia Elgoff, Mary A. Rutherford, Jonathan O’Muircheartaigh, Raphael Tomi-Tricot, Shaihan Malik, Lucillio Cordero-Grande, Joseph V. Hajnal, Serena J. Counsell

## Abstract

**Purpose:** To validate a retrospective motion correction technique, Distributed and Incoherent Sample Orders for Reconstruction Deblurring using Encoding Redundancy (DISORDER), for pediatric brain morphometry.

**Methods:** Two T1-weighted MPRAGE 3D datasets were acquired at 3T in thirty-seven children, median age 7.75 years; one with conventional linear phase encoding and one using DISORDER. MPRAGE images were scored as motion-free or motion-corrupt.

Cortical morphometry and regional brain volumes were measured with FreeSurfer, subcortical grey matter (GM) with FSL-FIRST, and hippocampal volumes with HippUnfold. Intraclass correlation coefficient (ICC) was used to determine agreement between the 2 MPRAGE acquisitions. Mann-Whitney *U* was used to test the difference between measures obtained using DISORDER and (i) motion-free and (ii) motion-corrupt conventional MPRAGE data.

**Results:** ICC measures were good/excellent for most subcortical GM (motion-free, 0.75-0.96; motion-corrupt, 0.62-0.98) and regional brain volumes (motion-free 0.47-0.99; motion-corrupt, 0.54-0.99) between conventional MPRAGE and DISORDER data, except for the amygdala and nucleus accumbens (motion-free, 0.38-0.65; motion- corrupt, 0.1-0.42). However, these values were less consistent for motion-corrupt conventional MPRAGE data for hippocampal volumes (motion-free 0.65-0.99; motion-corrupt, 0.11-0.91) and cortical measures (motion-free 0.76-0.98; motion- corrupt, 0.09-0.74). Mann-Whitney *U* showed percentage differences in measures obtained with motion-corrupt conventional MPRAGE compared to DISORDER data were significantly greater than in those obtained using motion-free conventional MPRAGE data in 22/58 structures.

**Conclusion:** In the absence of motion, morphometric measures obtained using DISORDER are largely consistent with those from conventional MPRAGE data, whereas improved reliability is obtained by DISORDER for motion-degraded scans. This study validates DISORDER for brain morphometric studies in children.

## Introduction

Structural magnetic resonance imaging (MRI) provides detailed information about brain morphometry. However, obtaining high-quality MR images in children can be challenging. In a clinical setting, good-quality MRI data is essential for accurate radiological assessment and in research, high-quality imaging data is required to assess morphometry. Head motion in particular is a common cause of image degradation in pediatric research settings and motion-induced artifacts can significantly influence estimates of brain metrics derived from MRI [1–3].

Multiple strategies are available for correcting intra-scan motion, including prospective and retrospective methods. Prospective methods track data describing the head pose, which is used to update the acquisition in real-time to ensure the field-of-view (FOV) remains aligned with the moving head [4]. These approaches are applicable across a wide range of sequences and scanners, however, they may require additional hardware and scanner modifications [4]. Retrospective motion correction techniques first collect k-space data and then correct for the effects of motion following acquisition, during the reconstruction stage. In a pediatric cohort, a retrospective technique improved T1-weighted image quality and automated segmentation of cortical and subcortical structures compared to a prospective motion correction technique [5]. Retrospective motion correction techniques include Distributed and Incoherent Sample Orders for Reconstruction Deblurring by using Encoding Redundancy (DISORDER), a sampling re-ordering scheme in the phase encoding plane [6, 7] which has been shown to improve image quality in a pediatric cohort [8]. However, brain morphometric measures obtained from DISORDER data have not been evaluated.

In addition to the sequence used to acquire the images, the choice of software for automated segmentation of brain structures needs to be considered. Several software packages are available for MRI structural analyses, including FreeSurfer [9], and FSL-FIRST [10]. Both of these segmentation tools have been used in cortical and subcortical GM morphometric analyses in pediatric studies [11–17]. Previous work has suggested that FreeSurfer performs best in cortical analyses [14], and FSL-FIRST is closer to manual segmentation in subcortical grey matter (GM) analyses in pediatric populations [12]. HippUnfold is a recently developed hippocampal-specific segmentation approach [18] and has been used previously to delineate hippocampal subfields in a pediatric population [19].

The aim of this study was to evaluate the use of DISORDER [6] in children aged 7-8 years. Conventional linear phase encoding and DISORDER Magnetization Prepared Rapid Gradient-Echo (MPRAGE) acquisitions were acquired. Brain morphometric measures were obtained using FreeSurfer [9] FSL-FIRST [10] and HippUnfold [18] and these measures were compared between the two MPRAGE acquisitions.

## Materials and Methods

### Participants

Data for this study were acquired as part of the ICONIC (Impact of Congenital Heart Disease on Neurodevelopment in Childhood) study between August 2022 and August 2023. Research ethics committee approval was granted (Ref: 22/WA/0014). Informed, written parental consent was obtained and written assent was obtained from participants.

### Magnetic resonance imaging acquisition

MRI was performed on a 3T scanner (MAGNETOM Vida, Siemens Healthcare, Erlangen, Germany) located at St Thomas’ Hospital (London, UK). The children were asked to stay still during scanning while watching a movie of their choice. Two T1 weighted MPRAGE 3D image datasets were acquired in the sagittal plane; one with conventional linear phase encoding (referred to as conventional MPRAGE) (TR = 2200 ms, TE = 2.46 ms, flip angle = 8°, FOV = 204 x 224 x 176, voxel size = 1.1 x 1.07 x 1.07 mm^3^, acceleration factor = 2) and one using the DISORDER scheme (TR = 2200 ms, TE = 2.45 ms, flip angle = 8°, FOV = 210 x 224 x 176, voxel size = 1.1 x 1.07 x 1.07 mm^3^, no acceleration). T2-weighted turbo spin echo, diffusion-weighted imaging, fluid attenuated inversion recovery imaging, and resting-state functional MRI were also acquired but not analysed as part of this study.

### Motion correction

Rigid-body motion correction ability is sensitive to the *k*-space encoding order [6, 7]. Combined with the aligned sensitivity encoding (SENSE) framework, which achieves intrascan motion estimation and correction by dividing k-space samples and accounting for different motion states in each temporal segment [7], DISORDER aims to improve motion tolerance of volumetric structural brain MRI by ensuring that the acquisition of each shot of *k*-space contains a series of samples distributed incoherently throughout *k*-space [6]. Since every shot contains samples distributed in k-space, low and high resolution information is available for motion estimation and chances are reduced for large uncovered k-space areas after a given rotation compensation is applied in reconstruction [6]. These significantly increase the ability to estimate head pose and obtain high-quality motion corrected reconstructions in the presence of rigid motion [6].

For reconstruction, the motion parameters and the image are jointly estimated. Starting from a reconstruction assuming no motion, a first approximation of the motion parameters for each *k*-space shot is estimated. Then a new volume is reconstructed with the motion parameters, and the method alternates between motion estimation and reconstruction until convergence [6, 7]. DISORDER reconstruction was performed using MATLAB R2018b (MathWorks, Inc., Natick, Massachusetts), and implemented based on its open-source implementation (available at https://github.com/mriphysics/DISORDER). Fig. 1 shows MPRAGE data acquisitions from two subjects: (A, D) conventional MPRAGE, (B, E) uncorrected and (C, F) motion corrected MPRAGE acquired with the DISORDER scheme.

**Fig. 1.**
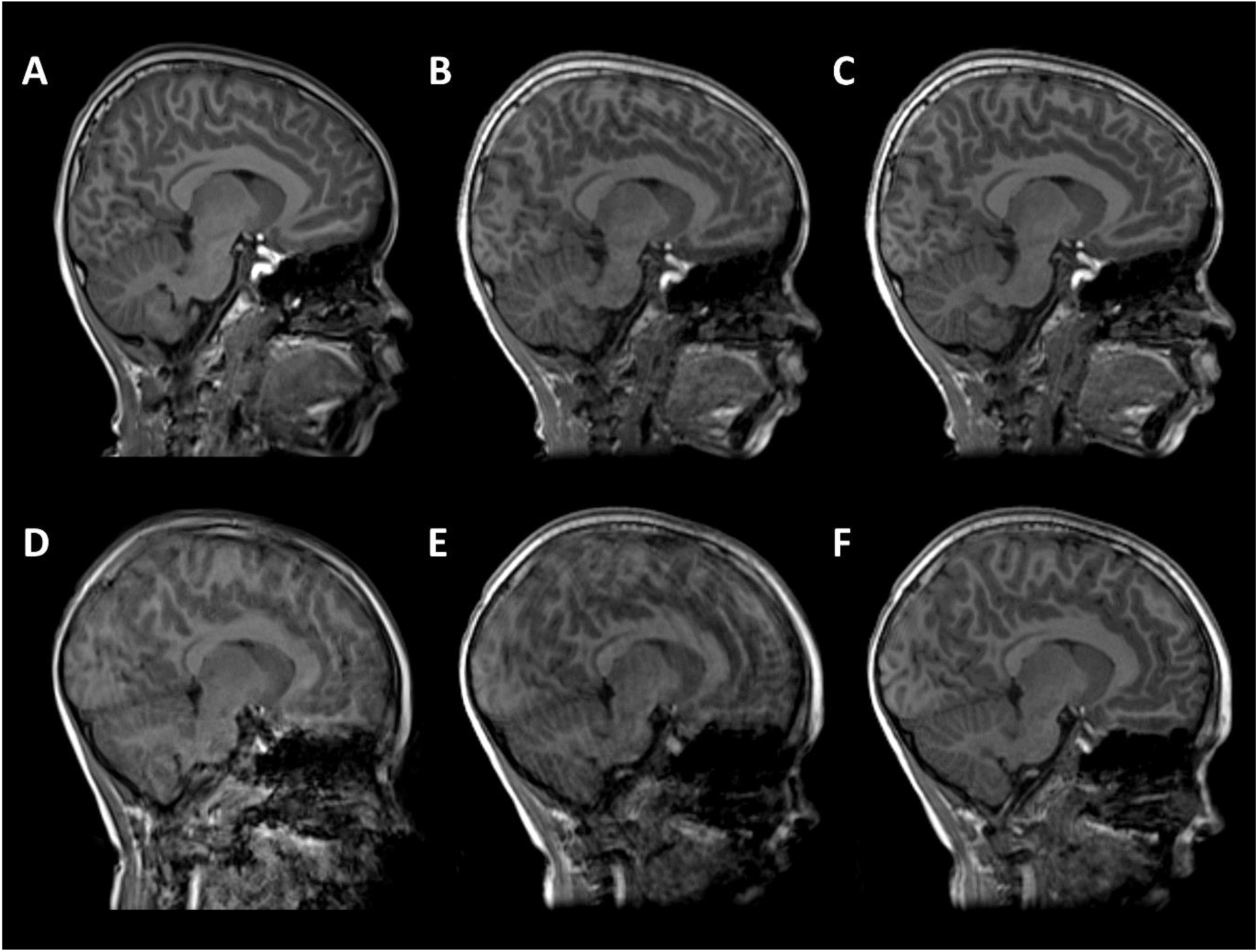
T1-weighted MPRAGE images from two subjects. (A-C) male, 7 years 11 months with good quality conventional MPRAGE data; (D- F) male,7 years 9 months with motion-corrupt conventional MPRAGE data. (A, D) conventional MPRAGE (B, E) DISORDER before motion correction and (C, F) DISORDER after motion correction.

### Image quality evaluation

Image quality was visually inspected by a pediatric neuroradiologist with more than 10 years of experience and an experienced image analyst. Sagittal, coronal and axial views of the MPRAGE data were presented to the raters in a randomized and blinded fashion and were scored as either no evidence of motion (referred to as motion-free) or motion-corrupt.

### Image preprocessing

After motion correction, the MPRAGE data underwent intensity normalization using ANTs [20]. Removal of Gibbs ringing artefacts was performed using a 3D extension to the unringing method proposed by Kellner and colleagues [21, 22]. Following bias field correction and unringing, images were preprocessed using the Human Connectome Project (HCP) minimal preprocessing pipeline [23].

MPRAGE data were aligned to MNI space using rigid registration [23]. After this alignment, an initial brain extraction is performed using linear and non-linear registration of the image to the MNI template [23]. The MPRAGE data were skull-stripped using SynthStrip, a deep learning-based brain extraction tool [24]. Using an optimised version of FreeSurfer’s recon-all pipeline (FreeSurfer v7.3.2, (https://surfer.nmr.mgh.harvard.edu/) [23, 25, 26], brain tissue was segmented into white matter (WM), grey matter (GM) and cerebrospinal fluid (CSF), and white and pial cortical surfaces were reconstructed [25].

### Metrics of interest

Subcortical GM volumes (thalamus, caudate nucleus, putamen, globus pallidus, amygdala, nucleus accumbens and brainstem) were segmented and volumes extracted using FSL-FIRST v6.0.5.2 [10].

Total hippocampal and hippocampal subfield volumes were segmented and volumes extracted using HippUnfold [18]. HippUnfold uses a U-Net neural network architecture to segment hippocampal GM and structures surrounding the hippocampus. Volumetric subfields were generated by filling the voxels between inner and outer surfaces with the corresponding subfield labels [18]. Subfield segmentations include the cornu ammonis (CA1, CA2, CA3, CA4), subiculum, dentate gyrus (DG) and the stratum radiatum lacunosum and moleculare (SRLM) (Fig. 2).

**Fig. 2.**
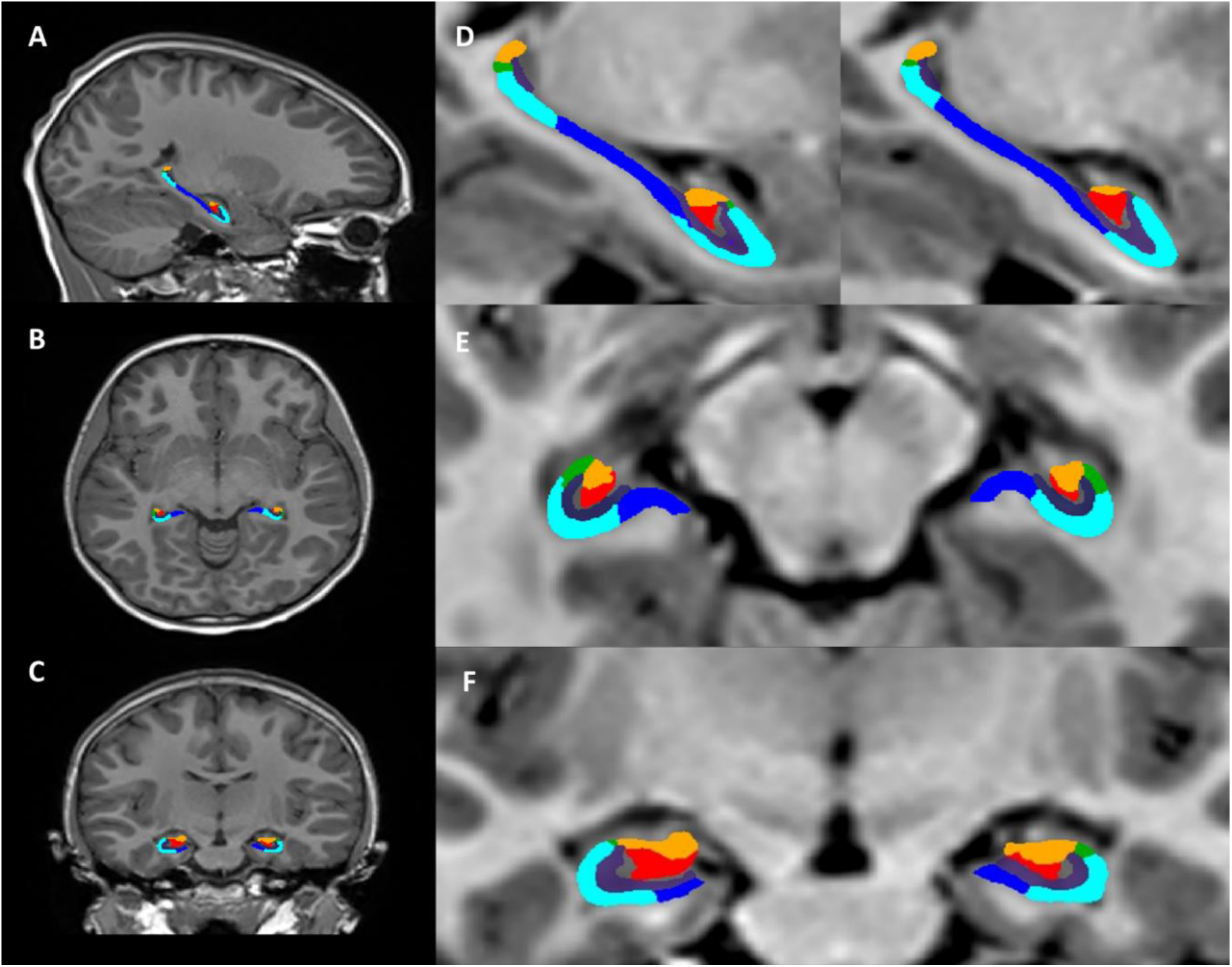
Hippocampal segmentation from one representative subject (female, 7 years 11 months). Segmentation of the hippocampus using HippUnfold in the sagittal (A, D), axial (B, E), and coronal planes (C, F). Hippocampal subfields; subiculum (dark blue), CA1 (light blue), CA2 (green), CA3 (orange), CA4 (red), SRLM (purple).

Volumes and surfaces generated using FreeSurfer were converted to standard formats used by Connectome Workbench, and a volume segmentation of the cortical ribbon was generated [23]. WM, ventral diencephalon, corpus callosum (anterior, mid-anterior, central, mid-posterior, posterior), lateral ventricles, third ventricle, fourth ventricle, cerebellar cortex, cerebellar WM and total cerebellum (cerebellar cortex + cerebellar WM) volumes were extracted using FreeSurfer.

Cortical GM measures (cortical surface area (SA), cortical GM volume, cortical thickness (CT), mean curvature, cortical gyrification) were also extracted using FreeSurfer. Cortical parcellations were computed based on the Desikan-Killiany atlas [27]. Cortical thickness was calculated as the average distance between the GM/WM boundary and the pial surface at each vertex within each region of interest (ROI) [25]. The surface area was calculated on the pial surface as the sum of the areas of each tessellation falling within a given ROI [25]. Mean curvature is calculated as 1/r, where r is the radius of an inscribed circle and mean curvature is the average of the two principal curvatures. Global measures of CT, surface area, mean curvature and GM volume were calculated as averages or sum totals of these ROIs. To measure cortical gyrification, a local gyrification index value (lGI) was computed for every vertex on the pial surface of the brain [28, 29]. The lGI at each vertex of the reconstructed cortical surface was measured as the ratio between the smoothed outer perimeter and inner buried contour of the cortex resulting in individual gyrification maps [28, 29]. The average value of all the individual vertex lGI values was used to calculate the mean lGI for each hemisphere [28, 29].

### Visual inspection of segmentations

Prior to statistical analysis, all segmentations were manually inspected using Freeview to ensure the accuracy of the cortical ribbon and tissue segmentation. Manual corrections were performed if delineation of the pial surface and WM boundary was poor, with defects spanning multiple slices. Edits were made according to instructions provided by FreeSurfer developers, (https://surfer.nmr.mgh.harvard.edu/fswiki/FsTutorial/TroubleshootingData).

### Statistical analysis

Statistical analyses were performed and graphs were generated using R v4.2.2 (https://www.r-project.org/) and R-Studio (https://rstudio.com/). Data were assessed for normality using Shapiro–Wilk tests, skewness, and kurtosis values. Descriptive statistics are shown as mean and standard deviation for normally distributed data and median and range for variables with a skewed distribution. False discovery rate (FDR) was used to correct for multiple comparisons and pFDR < 0.05 was considered significant [30].

### Intraclass correlation coefficients

Intraclass correlation coefficient (ICC) is an established measure of agreement between raters/measures. Here we calculated ICC as a measure of agreement for all morphological measures extracted from both MPRAGE acquisitions. The ICC is calculated as the proportion of overall variance that is explained by between-subject variance. ICC values greater than 0.90 indicate excellent reliability, values between 0.75 and 0.9 indicate good reliability, values between 0.5 and 0.75 indicate moderate reliability and values less than 0.5 are indicative of poor reliability [31].

### Percentage difference between brain morphometric measures

In order to assess the difference between morphometric measures obtained using DISORDER and (i) motion-free and (ii) motion-corrupt conventional MPRAGE images, the percentage difference between measures was computed using the formula: Percentage difference = [(DISORDER – conventional MPRAGE)/conventional MPRAGE] ∗ 100 [16]. Negative values indicate an underestimation of measures obtained with DISORDER compared to the conventional MPRAGE, while positive values indicate an overestimation of measures obtained with DISORDER relative to conventional MPRAGE data. Mann-Whitney *U* was used to test the effect of motion on the percentage difference.

## Results

### Participants

Sixty children were enrolled in the ICONIC study between August 2022 and August 2023, including 12 children with congenital heart disease (CHD) and 48 healthy controls. Twenty-three children were excluded from the analysis; conventional MPRAGE data not acquired (n=20), MPRAGE with DISORDER data not acquired (n=3). The final analysis included 37 children who underwent MR imaging at a median (range) age of 7.75 (7.5 – 8.33) years (Table 1); 20 children with motion-free conventional MPRAGE data and 17 children with motion-corrupt conventional MPRAGE data.

**Table 1.** Demographics of the study population.

|  | Motion-free<br>conventional MPRAGE<br>(N=20) | Motion-corrupt<br>conventional MPRAGE<br>(N=17) | Total<br>(N=37) |
| --- | --- | --- | --- |
| Median age at<br>scan [range],<br>(years) | 7.75<br>[7.5 - 8.17] | 7.83<br>[7.58 – 8.33] | 7.75<br>[7.5 - 8.33] |
| Male, no (%) | 10 (50) | 10 (59) | 20 (54) |

Out of the 17 children with motion-corrupt conventional MPRAGE data, 14 children had motion-free DISORDER data and 3 children had DISORDER data with minimal motion following reconstruction. All 20 children with motion-free conventional MPRAGE data had motion-free DISORDER data following reconstruction.

FreeSurfer’s reconstruction and segmentation failed in 1 child with motion-corrupt conventional MPRAGE data however, FSL-FIRST and HippUnfold were able to segment subcortical GM structures and hippocampal subfields respectively in this participant.

Example conventional MPRAGE and DISORDER images with cortical segmentation overlays are shown in Fig. 3 for a subject with motion-free conventional MPRAGE and a subject with motion-corrupt conventional MPRAGE data (Fig. 3).

**Fig. 3.**
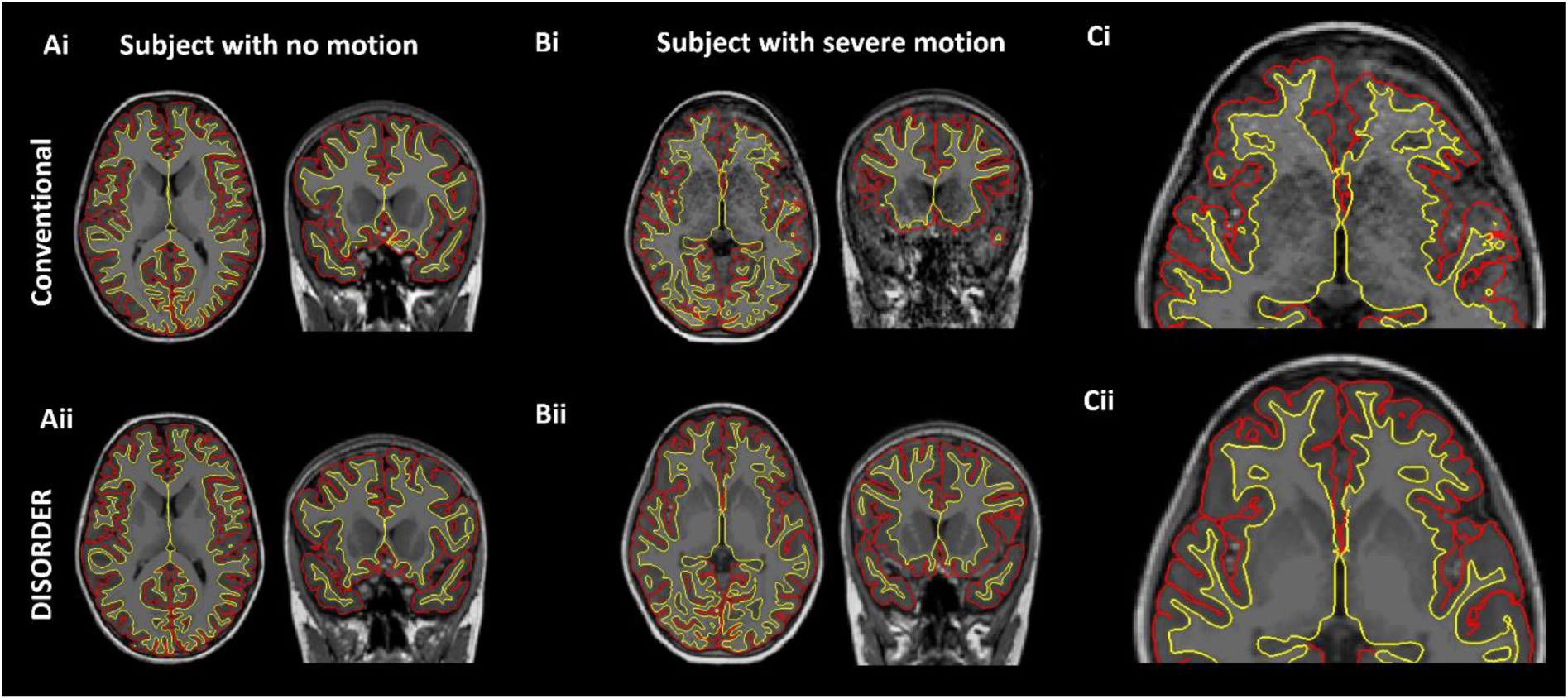
Cortical segmentations. The top row shows conventional MPRAGE acquisition and the bottom row shows the DISORDER MPRAGE acquisition. (Ai and ii) Female, 7 years 8 months no head motion. (B and C i and ii) Female, 8 years with head motion. (Ci) Motion corrupt conventional MPRAGE data created incorrect pial (red) and white matter (yellow) boundaries while DISORDER data (Cii) resulted in accurate cortical segmentations.

### Results of analyses comparing conventional MPRAGE and DISORDER data

#### Intraclass correlation coefficients

##### Subcortical grey matter volumes

ICC measures between motion-free conventional MPRAGE data and DISORDER data were excellent for the bilateral thalamus, bilateral caudate nucleus and brainstem; good for the bilateral putamen and globus pallidus; moderate for the right amygdala and left nucleus accumbens; and poor for the left amygdala and right nucleus accumbens (Table 2).

**Table 2.**
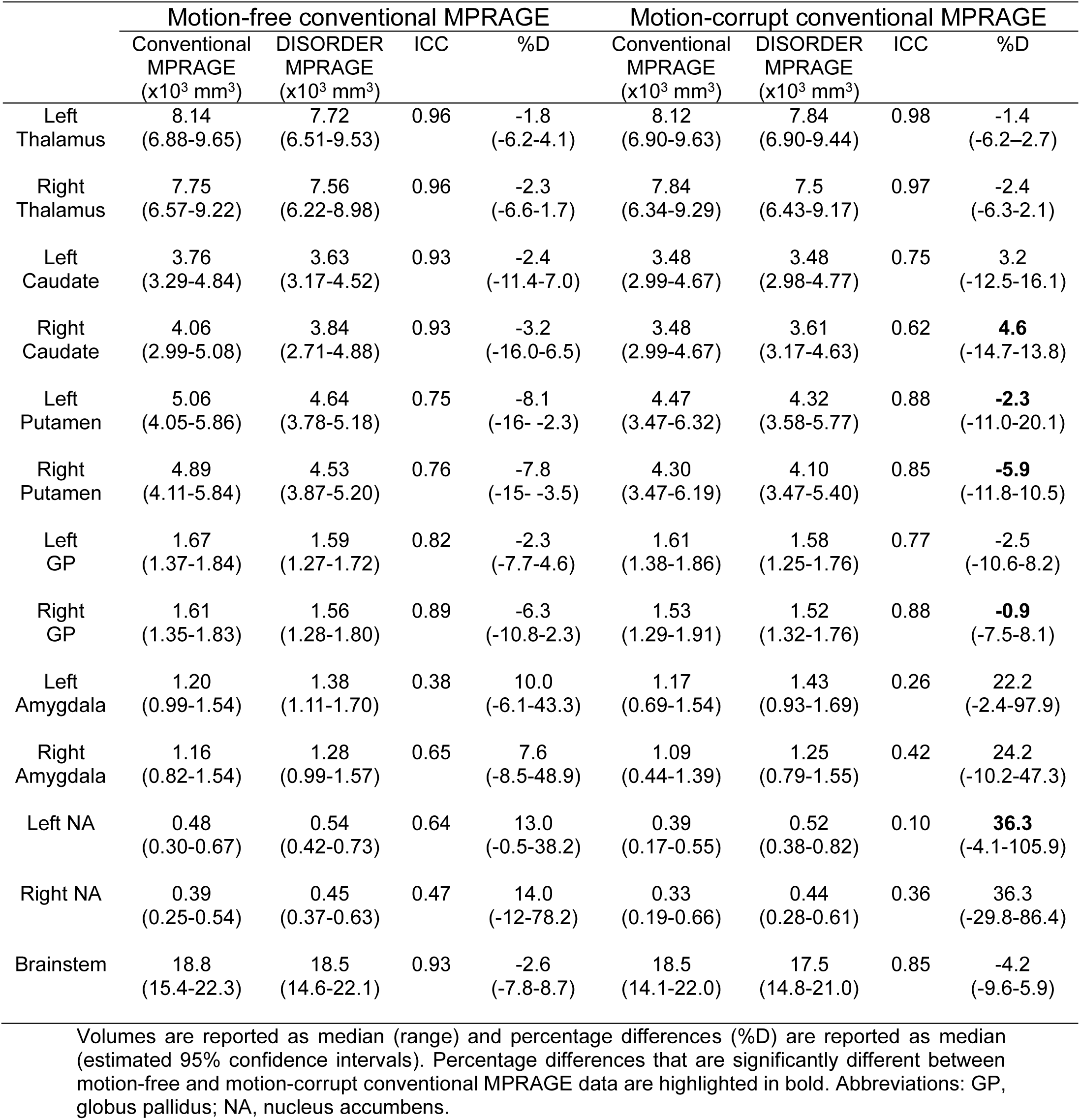
ICC and percentage differences between subcortical grey matter volumes obtained with conventional MPRAGE and DISORDER data.

ICC measures between motion-corrupt conventional MPRAGE and DISORDER data were excellent for the thalamus bilaterally; good for the bilateral putamen, bilateral globus pallidus, left caudate nucleus and brainstem; moderate for the right caudate nucleus; and poor for the bilateral amygdala and nucleus accumbens (Table 2).

##### Hippocampal volumes

ICC measures between motion-free conventional MPRAGE data and DISORDER data were excellent for the bilateral CA1, right CA3, bilateral CA4, bilateral dentate gyrus, bilateral SRLM and bilateral total hippocampal volume; good for the right subiculum, left CA3 and right CA2; and moderate for the left subiculum and left CA2 (Table 3).

**Table 3.** ICC and percentage differences between hippocampal volumes obtained with conventional MPRAGE and DISORDER data.

|  | Motion-free conventional MPRAGE |  |  |  | Motion-corrupt conventional MPRAGE |  |  |  |
| --- | --- | --- | --- | --- | --- | --- | --- | --- |
|  | Conventional MPRAGE (x10<sup>3</sup> mm<sup>3</sup>) | DISORDER MPRAGE (x10<sup>3</sup> mm<sup>3</sup>) | ICC | %D | Conventional MPRAGE (x10<sup>3</sup> mm<sup>3</sup>) | DISORDER MPRAGE (x10<sup>3</sup> mm<sup>3</sup>) | ICC | %D |
| Left subiculum | 0.52<br>(0.48-0.68) | 0.51<br>(0.44-0.68) | 0.74 | -6.6<br>(-18.1-7.2) | 0.47<br>(0.19-0.69) | 0.47<br>(0.41-0.66) | 0.17 | -3.6<br>(-20.6-95.4) |
| Right subiculum | 0.55<br>(0.40-0.66) | 0.51<br>(0.39-0.58) | 0.81 | -6.0<br>(-19.0-1.6) | 0.53<br>(0.34-0.74) | 0.50<br>(0.38-0.65) | 0.31 | -2.0<br>(-31.0-29.2) |
| Left CA1 | 0.86<br>(0.63-1.15) | 0.88<br>(0.68-1.11) | 0.99 | 1.8<br>(-3.6-6.0) | 0.78<br>(0.49-1.14) | 0.83<br>(0.63-1.06) | 0.43 | 4.0<br>(-11.7-86.7) |
| Right CA1 | 0.93<br>(0.82-1.15) | 0.94<br>(0.81-1.12) | 0.96 | 0.7<br>(-3.2-7.1) | 0.84<br>(0.65-1.26) | 0.89<br>(0.70-1.20) | 0.91 | 2.9<br>(-8.3-13.6) |
| Left CA2 | 0.12<br>(0.08-0.16) | 0.11<br>(0.07-0.13) | 0.65 | -3.5<br>(-19.8-28.4) | 0.12<br>(0.04-0.34) | 0.11<br>(0.06-0.21) | 0.63 | -17.3<br>(-68.7-41.2) |
| Right CA2 | 0.10<br>(0.07-0.14) | 0.094<br>(0.07-0.12) | 0.82 | -6.4<br>(-26.3-19.9) | 0.079<br>(0.04-0.14) | 0.094<br>(0.06-0.12) | 0.90 | **14.0**<br>(-9.7-75.2) |
| Left CA3 | 0.32<br>(0.25-0.40) | 0.30<br>(0.21-0.38) | 0.86 | -6.9<br>(-15.9-10.1) | 0.32<br>(0.13-0.55) | 0.28<br>(0.18-0.37) | 0.11 | -5.8<br>(-60.3-30.1) |
| Right CA3 | 0.25<br>(0.18-0.33) | 0.25<br>(0.20-0.31) | 0.90 | -4.3<br>(-14.0-17.6) | 0.21<br>(0.17-0.29) | 0.24<br>(0.19-0.35) | 0.62 | **10.6**<br>(-8.3-35.3) |
| Left CA4 | 0.15<br>(0.09-0.25) | 0.14<br>(0.09-0.24) | 0.94 | -5.2<br>(-25.6-24.8) | 0.15<br>(0.04-0.29) | 0.15<br>(0.08-0.29) | 0.57 | -3.1<br>(-34-137.3) |
| Right CA4 | 0.19<br>(0.12-0.31) | 0.17<br>(0.12-0.29) | 0.95 | -7.0<br>(-18.5-2.6) | 0.20<br>(0.13-0.31) | 0.18<br>(0.12-0.27) | 0.23 | -4.7<br>(-36.2-38.8) |
| Left DG | 0.13<br>(0.10-0.17) | 0.13<br>(0.10-0.16) | 0.95 | -2.9<br>(-12.0-1.4) | 0.11<br>(0.05-0.14) | 0.13<br>(0.10-0.14) | 0.38 | **-0.5**<br>(-8.2-112.4) |
| Right DG | 0.12<br>(0.10-0.16) | 0.13<br>(0.10-0.16) | 0.95 | -1.3<br>(-11.2-6.3) | 0.10<br>(0.07-0.16) | 0.13<br>(0.10-0.15) | 0.29 | **10.1**<br>(-6.0-66.8) |
| Left SRLM | 0.58<br>(0.50-0.78) | 0.57<br>(0.49-0.70) | 0.96 | -1.8<br>(-9.2-2.3) | 0.51<br>(0.24-0.77) | 0.55<br>(0.45-0.66) | 0.45 | -0.6<br>(-11.1-103.) |
| Right SRLM | 0.61<br>(0.51-0.77) | 0.59<br>(0.50-0.73) | 0.96 | -2.7<br>(-7.3-3.5) | 0.54<br>(0.43-0.78) | 0.57<br>(0.49-0.76) | 0.84 | -2.2<br>(-8.4-23.4) |
| Left total hippocampus | 2.74<br>(2.30-3.54) | 2.65<br>(2.28-3.20) | 0.96 | -2.1<br>(-8.0-0.5) | 2.43<br>(1.32-3.38) | 2.52<br>(2.05-3.17) | 0.54 | -2.6<br>(-11.7-21.4) |
| Right total hippocampus | 2.73<br>(2.35-3.37) | 2.66<br>(2.33-3.18) | 0.96 | -2.3<br>(-6.1-0.6) | 2.53<br>(1.94-3.43) | 2.60<br>(2.24-3.44) | 0.88 | **0.3**<br>(-5.9-17.9) |

Volumes are reported as median (range) and percentage differences (%D) are reported as median (estimated 95% confidence-intervals). Percentage differences that are significantly different between motion-free and motion-corrupt conventional MPRAGE data are highlighted in bold. Abbreviations: CA, cornu ammonius; DG, dentate gyrus; SRLM, stratum radiatum lacunosum and moleculare.

ICC measures between motion-corrupt conventional MPRAGE and DISORDER data were excellent for the right CA1 and CA2; good for the right SRLM and right total hippocampus; moderate for the left CA2, right CA3, left CA4 and left total hippocampus; and poor for the bilateral subiculum, left CA1, left CA3, right CA4, bilateral dentate gyrus and left SRLM (Table 3).

##### Regional brain volumes

ICC measures between motion-free conventional MPRAGE and DISORDER data were excellent for the bilateral WM, right cerebellar cortex, bilateral total cerebellum, anterior, mid-anterior, central and posterior CC, lateral ventricles bilateral, third and fourth ventricle; good for the bilateral ventral diencephalon, left cerebellar cortex and mid-posterior CC; moderate for the left cerebellar WM; and weak for the right cerebellar WM (Supplementary Table 1).

ICC measures between motion-corrupt conventional MPRAGE and DISORDER data were excellent for bilateral total cerebellum, posterior CC, bilateral lateral ventricles, third and fourth ventricle; good for bilateral ventral diencephalon, mid-posterior, anterior and mid-anterior CC; moderate for bilateral WM, bilateral cerebellar cortex, and central CC; and poor for bilateral cerebellar WM (Supplementary Table 1).

##### Cortical metrics

ICC measures between motion-free conventional MPRAGE data and DISORDER data were good or excellent for all cortical measures (Table 4).

**Table 4.** ICC and percentage differences between cortical measures obtained using conventional MPRAGE and DISORDER data.

|  | Motion-free conventional MPAGE |  |  |  | Motion-corrupt conventional MPAGE |  |  |  |
| --- | --- | --- | --- | --- | --- | --- | --- | --- |
|  | Conventional MPAGE | DISORDER MPAGE | ICC | %D | Conventional MPAGE | DISORDER MPAGE | ICC | %D |
| Left SA<br>($\times 10^4$ mm<sup>2</sup>) | 9.16<br>(8.08-9.97) | 9.67<br>(8.44-10.68) | 0.78 | 5.0<br>(0.7-14) | 7.97<br>(6.17-9.88) | 9.62<br>(7.82-12.11) | 0.52 | **22.2**<br>(9.8-33.2) |
| Right SA<br>($\times 10^4$ mm<sup>2</sup>) | 9.12<br>(7.93-9.97) | 9.69<br>(8.34-10.57) | 0.81 | 5.3<br>(2.7-12) | 8.05<br>(5.96-9.81) | 9.56<br>(7.88-11.83) | 0.53 | **21.2**<br>(8.6-32.4) |
| Left GM<br>($\times 10^5$ mm<sup>3</sup>) | 2.98<br>(2.48-3.4) | 2.96<br>(2.45-3.60) | 0.98 | -0.5<br>(-4.2-5.7) | 2.77<br>(2.04-3.4) | 3.04<br>(2.39-3.80) | 0.73 | **9.0**<br>(-7.0-29.8) |
| Right GM<br>($\times 10^5$ mm<sup>3</sup>) | 2.97<br>(2.44-3.4) | 2.93<br>(2.42-3.54) | 0.98 | -0.2<br>(-3.9-4.2) | 2.70<br>(1.96-3.4) | 3.10<br>(2.39-3.80) | 0.74 | **11.2**<br>(-3.9-26.4) |
| Left CT<br>(mm) | 2.82<br>(2.63-2.93) | 2.75<br>(2.51-3.09) | 0.78 | -1.4<br>(-6.9-4.7) | 2.82<br>(2.65-3.11) | 2.72<br>(2.45-2.91) | 0.26 | -4.3<br>(-17.6-3.9) |
| Right CT<br>(mm) | 2.78<br>(2.62-2.94) | 2.72<br>(2.50-3.05) | 0.81 | -1.7<br>(-5.5-5.0) | 2.84<br>(2.65-3.08) | 2.70<br>(2.45-2.90) | 0.09 | -5.2<br>(-15.4-3.7) |
| Left mean curvature<br>(1/mm) | 0.13<br>(0.12-0.14) | 0.13<br>(0.12-0.14) | 0.77 | 2.4<br>(-4.6-6.7) | 0.15<br>(0.14-0.16) | 0.13<br>(0.13-0.14) | 0.12 | **-9.1**<br>(-16.2-0.5) |
| Right mean curvature<br>(1/mm) | 0.13<br>(0.12-0.14) | 0.13<br>(0.12-0.14) | 0.76 | 1.9<br>(-3.7-7.1) | 0.15<br>(0.14-0.16) | 0.13<br>(0.13-0.14) | 0.16 | **-10.1**<br>(-14.7-2.3) |
| Left IGI | 3.47<br>(3.26-3.65) | 3.48<br>(3.23-3.68) | 0.92 | -0.1<br>(-3.6-3.5) | 3.22<br>(3.08-3.56) | 3.47<br>(3.29-3.95) | 0.45 | **7.6**<br>(0.2-12.7) |
| Right IGI | 3.43<br>(3.29-3.68) | 3.43<br>(3.26-3.71) | 0.92 | 0.2<br>(-3.7-4.0) | 3.22<br>(2.97-3.48) | 3.45<br>(3.25-3.89) | 0.48 | **9.1**<br>(0.1-17.2) |

Measures are reported as median (range) and percentage differences (%D) are reported as median (estimated 95% confidence-intervals). Percentage differences that are significantly different between motion-free and motion-corrupt conventional MPAGE data are highlighted in bold. Abbreviations: SA, surface area; GM, grey matter volume; CT, cortical thickness; IGI, local gyrification index.

ICC measures between motion-corrupt conventional MPRAGE and DISORDER data for cortical metrics were moderate for SA and GM volume bilaterally, and poor for other metrics (bilateral CT, mean curvature, and lGI) (Table 4).

#### Percentage difference between brain morphometric measures

##### Subcortical grey matter volumes

The median percentage difference for subcortical GM volumes between DISORDER and motion-free conventional MPRAGE data ranged from -1.8% to 14% and for motion-corrupt conventional MPRAGE data ranged from -1.4% to 36.3%. Table 2 shows the percentage difference for all subcortical GM volumes. The percentage difference between measures obtained using DISORDER and motion-corrupt conventional MPRAGE data was significantly greater than those obtained using motion-free conventional MPARGE data for the bilateral putamen, right caudate nucleus, right globus pallidus and left nucleus accumbens. There was no significant effect of motion on the percentage difference for the volume of any other subcortical GM structure (Fig. 4).

**Fig. 4.**
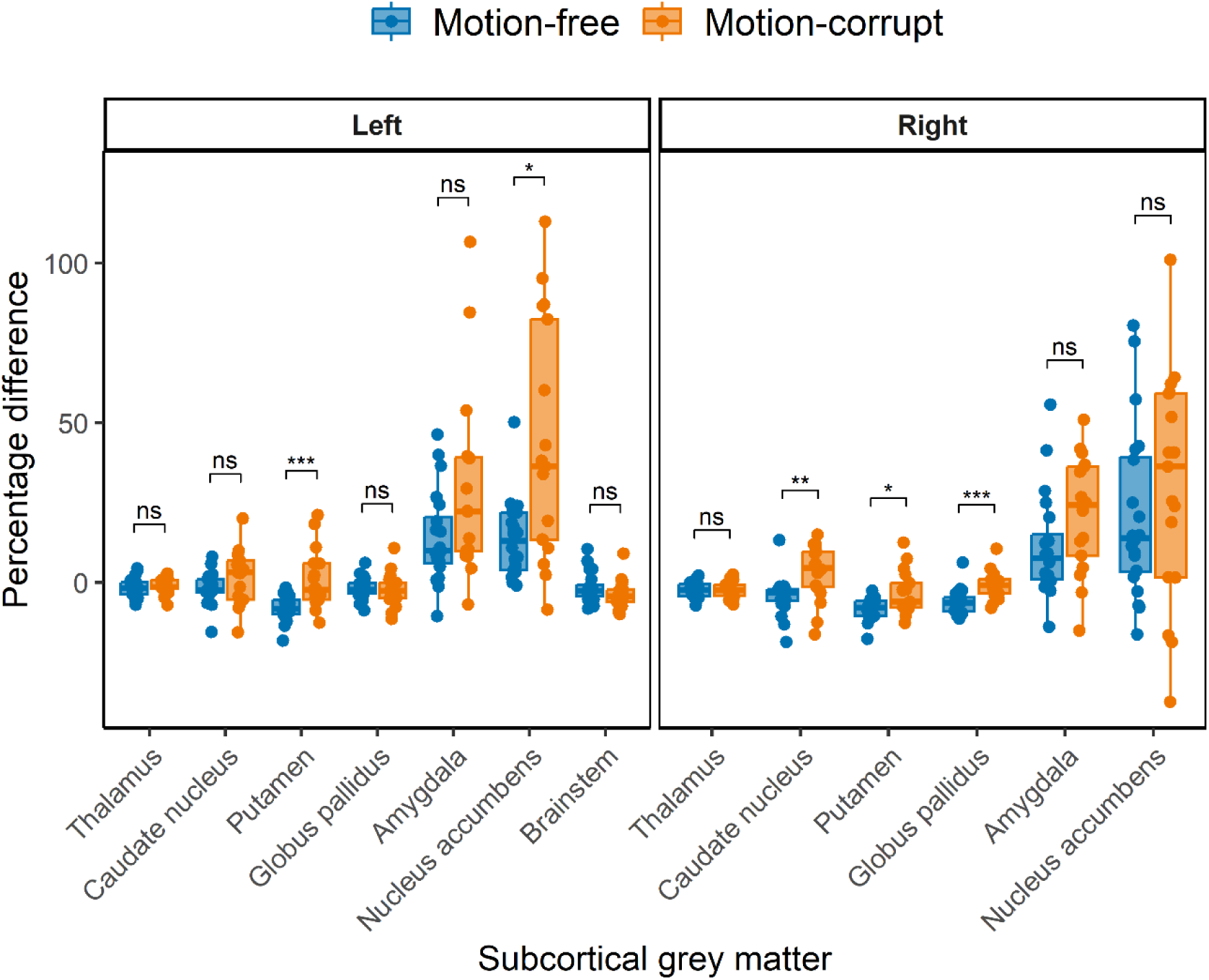
Percentage difference ((DISORDER – conventional MPRAGE/conventional MPRAGE) ∗ 100) between subcortical GM volumes obtained using DISORDER and motion-free and motion-corrupt conventional MPRAGE data. Results are shown separately for the left and right hemispheres (brainstem is included in the left panel). ns indicates a non-significant difference, * indicates a significant difference at the pFDR < 0.05 level, ** indicates a significant difference at the pFDR < 0.01 level, *** indicates a significant difference at the pFDR < 0.001 level.

##### Hippocampal volumes

The median percentage difference in hippocampal volumes between DISORDER and motion-free conventional MPRAGE data ranged from -7.0 % to 1.8% and for motion-corrupt conventional MPRAGE data ranged from -17.3% to 14%. Table 3 shows the percentage difference for all hippocampal volumes. The percentage difference between hippocampal volumes obtained using DISORDER and motion- corrupt conventional MPRAGE data was significantly greater than those obtained using motion-free conventional MPRAGE data for the right CA2, right CA3, bilateral dentate gyrus and right total hippocampus. There was no significant effect of motion on the percentage difference for the volume of any other hippocampal measure (Fig. 5).

**Fig. 5.**
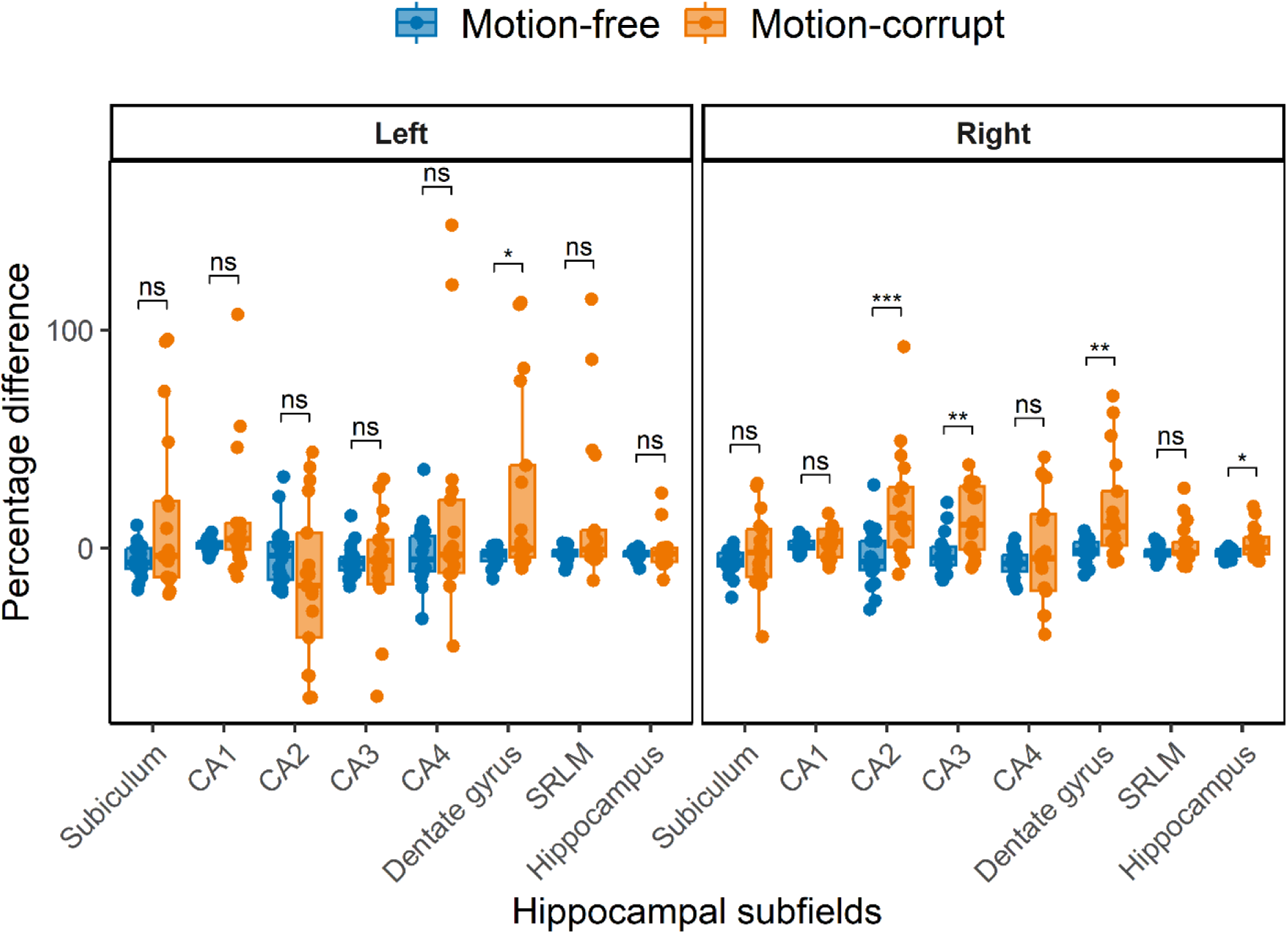
Percentage difference between hippocampal volumes obtained using DISORDER and motion-free and motion-corrupt conventional MPRAGE data. Results are shown separately for the left and right hemispheres. Abbreviations: CA, cornu ammonis; SRLM, stratum radiatum lacunosum and moleculare. ns indicates a non-significant difference, * indicates a significant difference at the pFDR < 0.05 level, ** indicates a significant difference at the pFDR < 0.01 level, *** indicates a significant difference at the pFDR < 0.001 level.

##### Regional brain volumes

The median percentage difference in regional brain volumes between DISORDER and motion-free conventional MPRAGE data ranged from -6.3% to 7.4% and for motion-corrupt conventional MPRAGE data ranged from -8.7% to 13%.

Supplementary Table 1 shows the percentage difference for all regional brain volumes. The percentage difference between measures obtained using DISORDER and motion-corrupt conventional MPRAGE data was significantly greater than those obtained using motion-free conventional MPRAGE data for the bilateral WM, left total cerebellum and mid-anterior corpus callosum volumes. There was no significant effect of motion on the percentage difference for any other regional brain volume (Supplementary Fig. 1).

##### Cortical metrics

The median percentage difference of regional brain volumes between DISORDER and motion-free conventional MPRAGE data ranged from -0.5% to 5.3% and for motion-corrupt conventional MPRAGE data ranged from -10.1% to 22.2%. Table 4 shows the percentage difference for all cortical metrics. The percentage difference between measures obtained using DISORDER and motion-corrupt conventional MPRAGE data was significantly greater than those obtained using motion-free conventional MPRAGE data for the bilateral SA, cortical GM volume, mean curvature and lGI. There was no significant effect of motion on the percentage difference for CT bilaterally (Fig. 6).

**Fig. 6.**
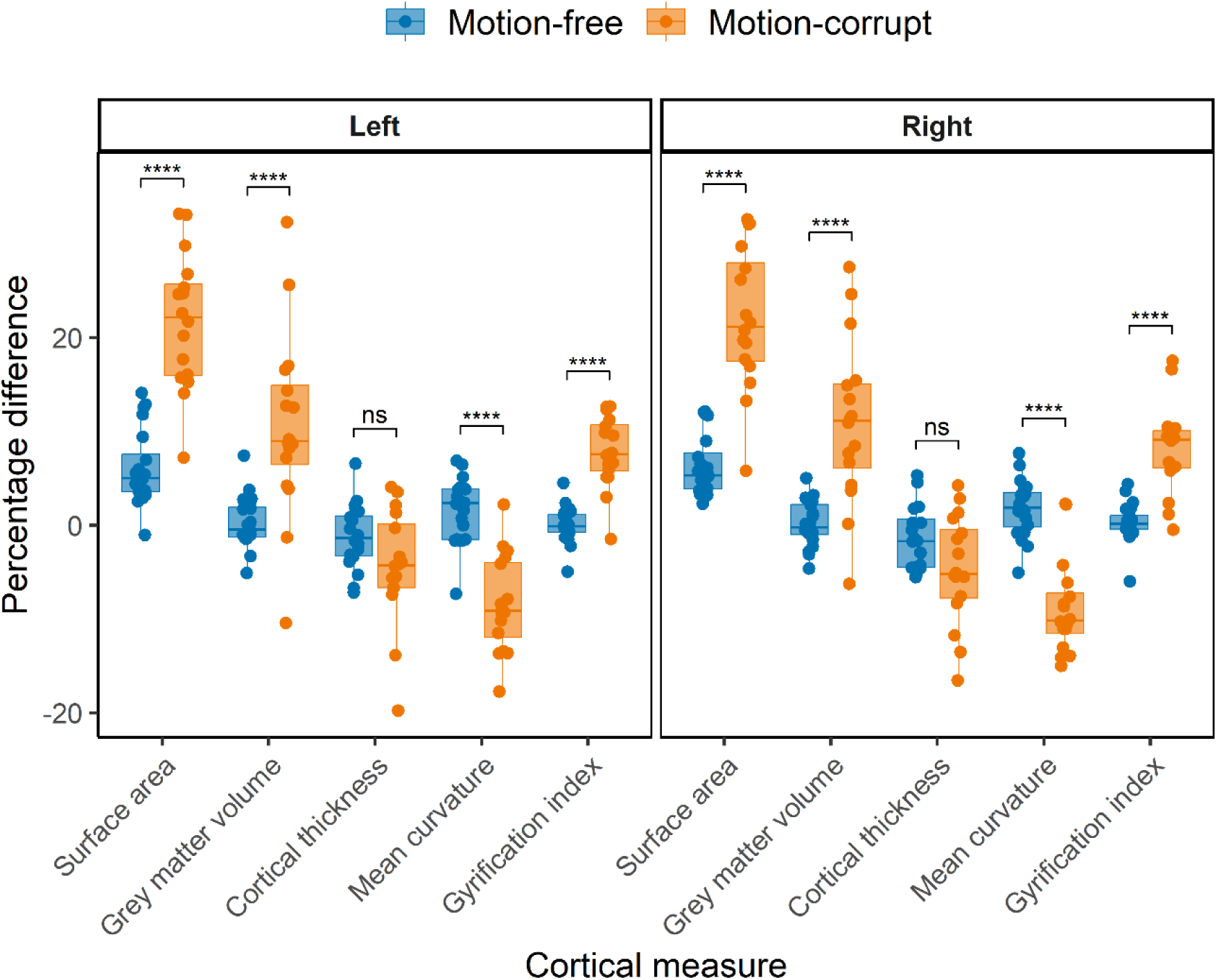
Percentage difference between cortical measures obtained using DISORDER and motion-free and motion-corrupt conventional MPRAGE data. Results are shown separately for the left and right hemispheres. ns indicates a non-significant difference, **** indicates a significant difference at the pFDR < 0.0001 level.

## Discussion

In this study, we evaluated the use of the DISORDER scheme for pediatric brain morphometry by comparing regional and subcortical GM brain volumes, hippocampal subfield volumes and cortical measures derived from conventional linear phase encoding MPRAGE images with those of MPRAGE acquired with the DISORDER sampling scheme in children aged between 7 and 8 years. We studied a wide range of brain structures to include regions that would be of interest to a wide range of pediatric neuroimaging studies. Our results suggest that, for most regions, brain morphometric measures obtained from DISORDER MPRAGE and motion-free conventional MPRAGE data are highly consistent. Brain morphometric measures obtained from motion-corrupt conventional MPRAGE and DISORDER data showed greater variance in concordance, particularly in hippocampal and cortical metrics.

Obtaining high-quality MR images in children is challenging, and head motion is a major cause of image degradation in pediatric neuroimaging. Previous work demonstrated that the DISORDER framework improved image quality in a pediatric cohort (age range 2-18 years) [8]. In our study, a high proportion of data acquired with the conventional acquisition had evidence of motion artifact (17 out of 37 children, 46%), highlighting the need for motion correction approaches in pediatric populations. Furthermore, in 14/17 (82%) children with motion-corrupt conventional MPRAGE data, image quality was good following a motion corrected DISORDER acquisition, and only minimal motion artefact was apparent in the remaining 3 datasets. These findings suggest that DISORDER may reduce the number of failed examinations in clinical settings, in addition to being a useful tool in research neuroimaging studies.

Our results show that cortical measures obtained using DISORDER and motion-free conventional MPRAGE acquisitions were highly consistent. However, we found that cortical measures showed poorer concordance between DISORDER and motion- corrupt conventional MPRAGE data. We also report a significant effect of motion, as assessed by the percentage difference between measures obtained using DISORDER and motion-free and motion-corrupt conventional MPRAGE data, on most cortical measures, with >20% difference in SA between DISORDER and motion-corrupt conventional MPRAGE data (compared to a percentage difference of 5% between DISORDER and motion-free conventional MPRAGE data in SA measures), highlighting the importance of correcting for head motion when obtaining cortical morphometric measures. Our findings are consistent with previous studies that demonstrated a reduction in estimates of cortical GM volume [1, 32] and an increase in estimates of mean curvature [32] with subject motion.

In addition to cortical measures, we found that most subcortical GM and regional brain volumes showed excellent or good agreement between motion-free conventional and DISORDER acquisitions. Interestingly, we also found that good-excellent agreement of some subcortical GM and regional brain volumes between motion-corrupt conventional MPRAGE and DISORDER data. However, several subcortical GM and regional brain volumes showed a significant effect of motion when comparing the percentage difference in volumes obtained from motion-free and motion-corrupt conventional MPRAGE to DISORDER data, whereas the percentage difference in subcortical GM and regional volumes obtained from motion-free conventional MPRAGE were similar to percentage differences reported in scan-rescan reliability studies in adults [33].

It is notable that agreement between DISORDER and conventional MPRAGE data for amygdala and nucleus accumbens volumes were poor-moderate for both motion-free and motion-corrupt data. Less reliable segmentations of the amygdala and nucleus accumbens have been demonstrated in pediatric test-retest studies [5] and in studies comparing automated segmentation to manual delineation [12], suggesting that the segmentation of these structures may be more susceptible to measurement error due to their small size [33]. The amygdala and nucleus accumbens also demonstrated large percentage volume differences (>20% and 36% respectively) between DISORDER and motion-corrupt conventional MPRAGE data. Of note, the percentage volume differences observed here between DISORDER and motion-free conventional MPRAGE data for the amygdala and nucleus accumbens (7-14%) are similar to those reported in a scan-rescan study in adults using FSL-FIRST [33]. The large differences in estimates of brain morphometry in motion-corrupt MRI data have consequences for pediatric neuroimaging research, particularly case-control studies where reported morphometric differences may be related to motion-related bias and not genuine biological differences between groups.

Several studies have highlighted altered hippocampal volumetry in pediatric clinical populations, including in children with epilepsy [34], CHD [35], and ADHD [36]. However, in addition to the amygdala and nucleus accumbens, the hippocampus can be challenging to segment due to the interindividual variability in hippocampal size and anatomy [37]. In this study, we showed that volumes of hippocampal subfields, were similar between motion-free conventional MPRAGE and DISORDER data. However, hippocampal volumes showed more variability between acquisitions for motion-corrupt MPRAGE data. Furthermore, the percentage difference between DISORDER and motion-corrupt volumes of some hippocampal structures was >10%. Our data highlight that good quality MR data is required for hippocampal subfield segmentation in order to assess differences in these structures between at risk groups and controls, and to investigate associations with neurodevelopmental outcomes.

It is important to note that cerebellar WM demonstrated weak to moderate agreement between both motion-free and motion-corrupt conventional MPRAGE and DISORDER data. Previous work assessing test-retest agreement of automated tools for cerebellar segmentation in adults reported lower ICC values and greater volume differences in FreeSurfer-derived cerebellar WM volumes compared to cerebellar GM volumes [38], highlighting that care should be taken when interpreting results in these regions. However, in our study, the total cerebellar (cerebellar GM + cerebellar WM) volume demonstrated excellent agreement between DISORDER and conventional MPRAGE data, suggesting that studies focusing on the whole cerebellum can reliably use measures derived from DISORDER data.

## Limitations

We studied a narrow age range, between 7 and 8 years. Future studies incorporating a wider age range could further validate the use of DISORDER for brain morphometric analyses.

## Conclusion

DISORDER enables quantitative structural MRI in difficult-to-image pediatric populations and will facilitate quantitative morphometry in research studies. Our results suggest that morphometric measures obtained using DISORDER retrospective motion correction are largely concordant with those obtained using motion-free conventional MPRAGE data. Motion corrected DISORDER improves the overall accuracy of most morphometric measures when compared to motion-corrupt conventional MPRAGE data. This study validates the use of DISORDER for brain morphometric studies in children.

