## Supplementary Materials for "Verifying the concordance between motion corrected and conventional MPRAGE for pediatric morphometric analysis"

**Supplementary Table 1.** ICC and percentage differences between regional brain volumes obtained using conventional MPRAGE and DISORDER data.

|  | Motion-free conventional MPRAGE | | | | Motion-corrupt conventional MPRAGE | | |  |
| --- | --- | --- | --- | --- | --- | --- | --- | --- |
|  | Conventional MPRAGE (x10^3^ mm^3^) | DISORDER  MPRAGE  (x10^3^ mm^3^) | ICC | %D | ConventionalMPRAGE (x10^3^ mm^3^) | DISORDER  MPRAGE  (x10^3^ mm^3^) | ICC | %D |
| Left WM | 210.5  (181.3-246) | 211.7  (186.7-252.6) | 0.96 | 2.1  (-2.5-8.5) | 187.2  (130.3–249) | 212.8  (157.1- 279) | 0.69 | **13.0**  (3.5-20.7) |
| Right WM | 208.5  (180-241.2) | 209.6  (185.9-247.2) | 0.96 | 2.5  (-1.7-7.5) | 187.8  (129 -245.8) | 213.6  (156.6- 272.8) | 0.71 | **13.0**  (2.9-20.2) |
| Left ventral DC | 3.8  (3.1 – 4.4) | 3.7  (3.1 – 4.3) | 0.85 | -6.3  (-10.5-2.5) | 3.7  (3.0 – 4.5) | 3.5  (3.0 – 4.3) | 0.78 | -4.7  (-13-5.6) |
| Right ventral DC | 3.8  (3.2 – 4.7) | 3.6  (3.0 – 4.1) | 0.87 | -5.4  (-10.6-3.3) | 3.6  (3.1-4.5) | 3.5  (2.9 – 4.2) | 0.72 | -5.9  (-15-5.9) |
| Left cerebellar cortex | 56.6  (48.7-73.9) | 53.8  (45.7 – 67.6) | 0.85 | -3.9  (-19.5-2.2) | 59.2  (53.0 – 68.9) | 55.1  (49.5 –69.4) | 0.54 | -3.4  (-15-4.7) |
| Right cerebellar cortex | 56.3  (49.3–72.6) | 54.1  (46.2 – 72.3) | 0.92 | -2.0  (-14.8-4.3) | 58.9  (52.3 – 69.4) | 58.6  (54.1 –71.1) | 0.69 | 0.4  (-8.3-6.8) |
| Left cerebellar WM | 14.0  (11.5–21.8) | 14.6  (11.5 – 26.6) | 0.54 | 2.1  (-12.5-72.2) | 13.8  (10.5 – 17.8) | 16.3  (12.4 –24.3) | 0.19 | 7.7  (-4.4-67.9) |
| Right cerebellar WM | 13.2  (10.7–20.9) | 13.4  (10.7 – 23.1) | 0.47 | -0.5  (-27.7-66.6) | 13.8  (11.2 – 22.0) | 14.2  (11.5 –23.4) | 0.14 | -1.8  (-23.1-51.2) |
| Total left cerebellum | 70.4  (60.1-90.4) | 68.9  (57.7-93.1) | 0.95 | -2.6  (-8.8-3.0) | 73.4  (65.6-84.8) | 72.7  (66.4-86.1) | 0.94 | **0.6**  (-4.0-3.8) |
| Total right cerebellum | 71.1  (60.4-88.6) | 68.8  (57.7-89.0) | 0.95 | -2.1  (-6.2-3.1) | 74.5  (66.5-85.0) | 72.9  (65.7-86.8) | 0.90 | -0.3  (-5.4-5.0) |
| Posterior  CC | 0.84  (0.68–1.07) | 0.87  (0.73 – 1.07) | 0.92 | 1.5  (-4.5-23.3) | 0.8  (0.61 – 1.20) | 0.87  (0.54 – 1.20) | 0.96 | 1.1  (-8.1-16.1) |
| Mid-posterior CC | 0.51  (0.37-0.65) | 0.52  (0.42 -0.66) | 0.84 | 2.6  (-7.3-32.7) | 0.49  (0.39 – 0.66) | 0.50  (0.36 – 0.67) | 0.88 | 2.2  (-17.6-10.7) |
| Central  CC | 0.57  (0.36–0.97) | 0.61  (0.35 – 1.05) | 0.95 | 6.7  (-4.6-30.6) | 0.52  (0.34 – 0.71) | 0.54  (0.31 – 0.68) | 0.66 | -1.0  (-18.3-28.2) |
| Mid-anterior CC | 0.56  (0.35–0.85) | 0.62  (0.35 – 0.91) | 0.90 | 7.4  (-7.7-48.4) | 0.50  (0.34 – 0.86) | 0.49  (0.30 – 0.80) | 0.87 | **-8.7**  (-21.9-5.4) |
| Anterior  CC | 0.82  (0.65 – 1.1) | 0.84  (0.62 – 1.2) | 0.95 | 1.5  (-10-14.7) | 0.83  (0.63 – 1.40) | 0.85  (0.66 – 1.36) | 0.84 | 3.0  (-7.5-19.9) |
| Left LV  Mean (SD) | 4.6  (1.9) | 4.5  (1.9) | 0.99 | -2.9  (-14.5-4.8) | 6.22  (2.5) | 5.90  (2.6) | 0.99 | -3.6  (-21.8-0.7) |
| Right LV  Mean (SD) | 4.4  (1.80) | 4.4  (1.90) | 0.99 | 0.3  (-8.5-5.6) | 5.7  (3.12) | 5.5  (3.20) | 0.99 | -2.7  (-24.2-3.3) |
| Third ventricle  Mean (SD) | 0.79  (0.20) | 0.79  (0.20) | 0.99 | 1.0  (-8.3-9.7) | 0.87  (0.28) | 0.85  (0.25) | 0.97 | -2.7  (-13.4-8.2) |
| Fourth ventricle | 1.66  (1.12–2.90) | 1.74  (1.21 – 2.75) | 0.99 | 1.1  (-10.5-12.1) | 1.81  (1.0 – 3.09) | 1.79  (1.03 – 3.10) | 0.98 | 0.6  (-16.4-10.2) |

MPRAGE volumes are reported as median (range) unless stated otherwise, and percentage differences (%D) are reported as median (95% confidence interval). Percentage differences that are significantly different between motion-free and motion-corrupt MPRAGE data are highlighted in bold. Abbreviations: CC, corpus callosum; DC, diencephalon; LV, lateral ventricle; WM, white matter; SD, standard deviation.

**Supplementary Figure legends**

**Supplementary Fig. 1** Percentage difference between regional brain volumes obtained using DISORDER and motion-free and motion-corrupt conventional MPRAGE data.. ns indicates a non-significant difference, **** indicates a significant difference at the pFDR < 0.0001 level, *** indicates a significant difference at the pFDR < 0.001 level. Abbreviations: CC, corpus callosum.

**Supplementary Figures**

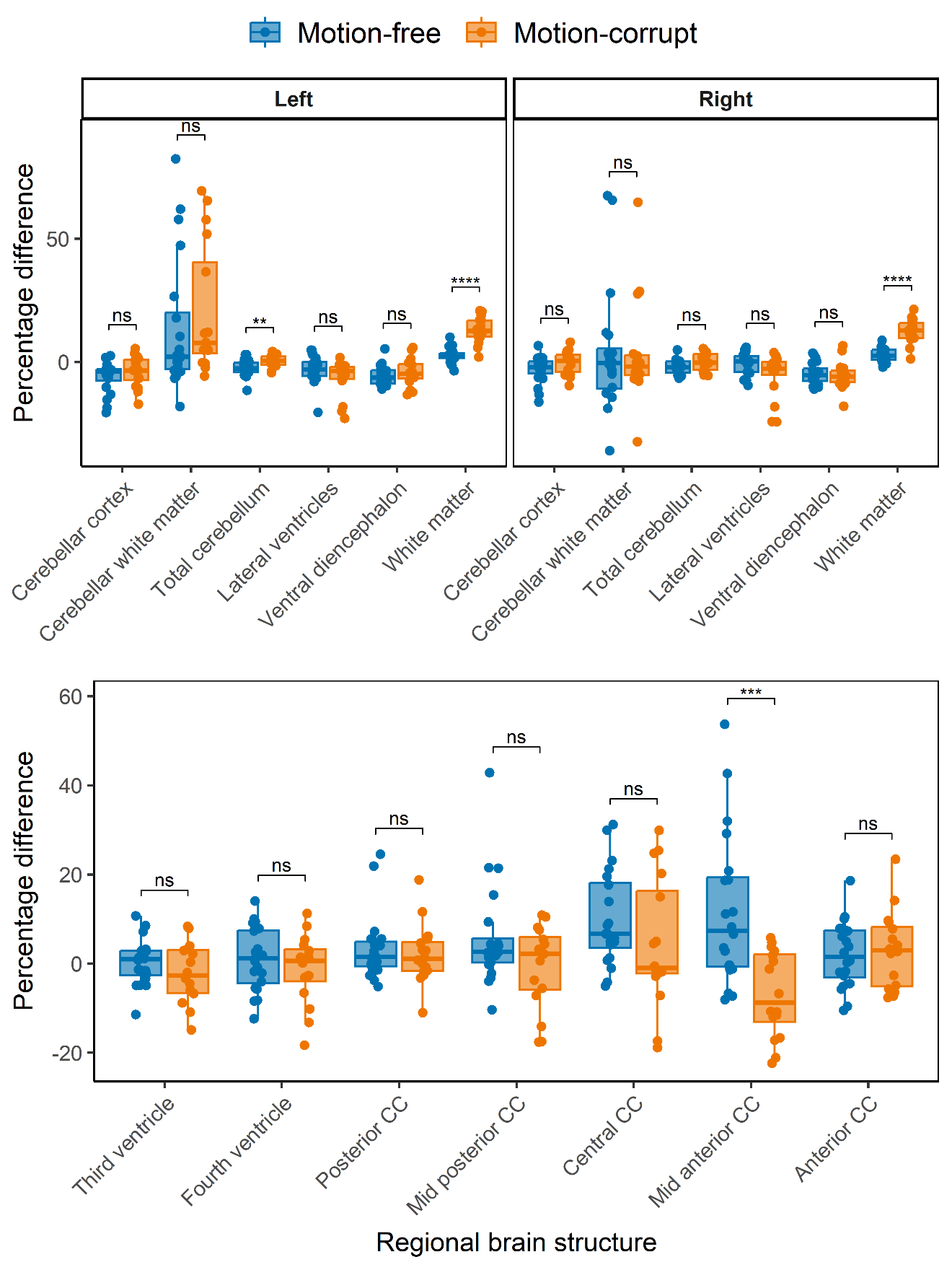

**Supplementary Fig. 1** Percentage difference between regional brain volumes obtained using DISORDER and motion-free and motion-corrupt conventional MPRAGE data.. ns indicates a non-significant difference, **** indicates a significant difference at the pFDR < 0.0001 level, *** indicates a significant difference at the pFDR < 0.001 level. Abbreviations: CC, corpus callosum.
